# Distinct timescales of RNA regulators enable the construction of a genetic pulse generator

**DOI:** 10.1101/377572

**Authors:** Alexandra Westbrook, Xun Tang, Ryan Marshall, Colin S. Maxwell, James Chappell, Deepak K. Agrawal, Mary J. Dunlop, Vincent Noireaux, Chase L. Beisel, Julius Lucks, Elisa Franco

**Affiliations:** Robert F. Smith School of Chemical and Biomolecular Engineering, Cornell University, Ithaca, NY 14853, United States; Department of Mechanical Engineering, University of California at Riverside, Riverside, CA 92521, United States; School of Physics and Astronomy, University of Minnesota, Minneapolis, MN 55455, United States; Department of Chemical and Biomolecular Engineering, North Carolina State University, Raleigh, NC 27695, United States; Department of Biosciences, Rice University, Houston, TX 77005, United States; Biomedical Engineering Department, Boston University, Boston, MA 02215, United States; Helmholtz Institute for RNA-based Infection Research (HIRI), Josef-Schneider-Str. 2 / D15, D-97080 Würzburg, Germany; Faculty of Medicine, University of Würzburg, Würzburg, Germany; Department of Chemical and Biological Engineering, Northwestern University, Evanston, IL 60208, United States

## Abstract

To build complex genetic networks with predictable behaviours, synthetic biologists use libraries of modular parts that can be characterized in isolation and assembled together to create programmable higher-order functions. Characterization experiments and computational models for gene regulatory parts operating in isolation are routinely employed to predict the dynamics of interconnected parts and guide the construction of new synthetic devices. Here, we individually characterize two modes of RNA-based transcriptional regulation, using small transcription activating RNAs (STARs) and CRISPR interference (CRISPRi), and show how their distinct regulatory timescales can be used to engineer a composed feedforward loop that creates a pulse of gene expression. We use a cell-free transcription-translation system (TXTL) to rapidly characterize the system, and we apply Bayesian inference to extract kinetic parameters for an ODE-based mechanistic model. We then demonstrate in simulation and verify with TXTL experiments that the simultaneous regulation of a single gene target with STARs and CRISPRi leads to a pulse of gene expression. Our results suggest the modularity of the two regulators in an integrated genetic circuit, and we anticipate that construction and modeling frameworks that can leverage this modularity will become increasingly important as synthetic circuits increase in complexity.

## INTRODUCTION

An important goal of synthetic biology is the development of rational methods for precise temporal control of gene expression, which is necessary to achieve sophisticated dynamic functions in engineered cells.^1^ Towards this broad goal, libraries of synthetic regulatory parts have been developed to give synthetic biologists control over distinct levels of gene expression.^2,3^ In order to create more complex networks, these parts need to be modular and composable,^4^ performing their function within the network with minimal undesired interactions. RNA provides a powerful platform to achieve this.

RNA-based regulators have become increasingly popular for building libraries of synthetic parts to orthogonally control many aspects of gene expression.^2,3,5-7^ RNA transcriptional regulators are particularly interesting because they can regulate RNA synthesis as a function of an RNA input and thus can be used to create genetic circuitry that propagates signals on the RNA level.^6,8^ These circuits have many potential advantages over protein-based circuits, including the ability to leverage RNA-folding algorithms and high-throughput structure determination to optimize regulatory part folding and function,^9^ reduced metabolic load for the host,^10^ and rapid signal propagation due to their fast degradation rates.^8^

Here we focus on building a simple genetic network by combining two modes of RNA-based transcriptional regulation: using small transcription activating RNAs (STARs)^11^ and clustered regularly interspaced short palindromic repeats (CRISPR) interference (CRISPRi).^12,13^ STARs activate gene expression through an interaction with a sequence specific target RNA. The target RNA resides in the 5’ UTR of the gene of interest and folds into a transcriptional terminator that halts transcription by causing the polymerase to fall off of the DNA complex before the downstream gene. When present, the activating RNA – called the STAR – binds to the target RNA to prevent terminator formation, thus allowing downstream transcription to turn gene expression ON (Figure 1A). Libraries of orthogonal STARs have been built and shown to work in many contexts, including within genomic DNA to reprogram cellular phenotypes, and to control multiple genes within a metabolic pathway.^2,11^

**Figure 1.**
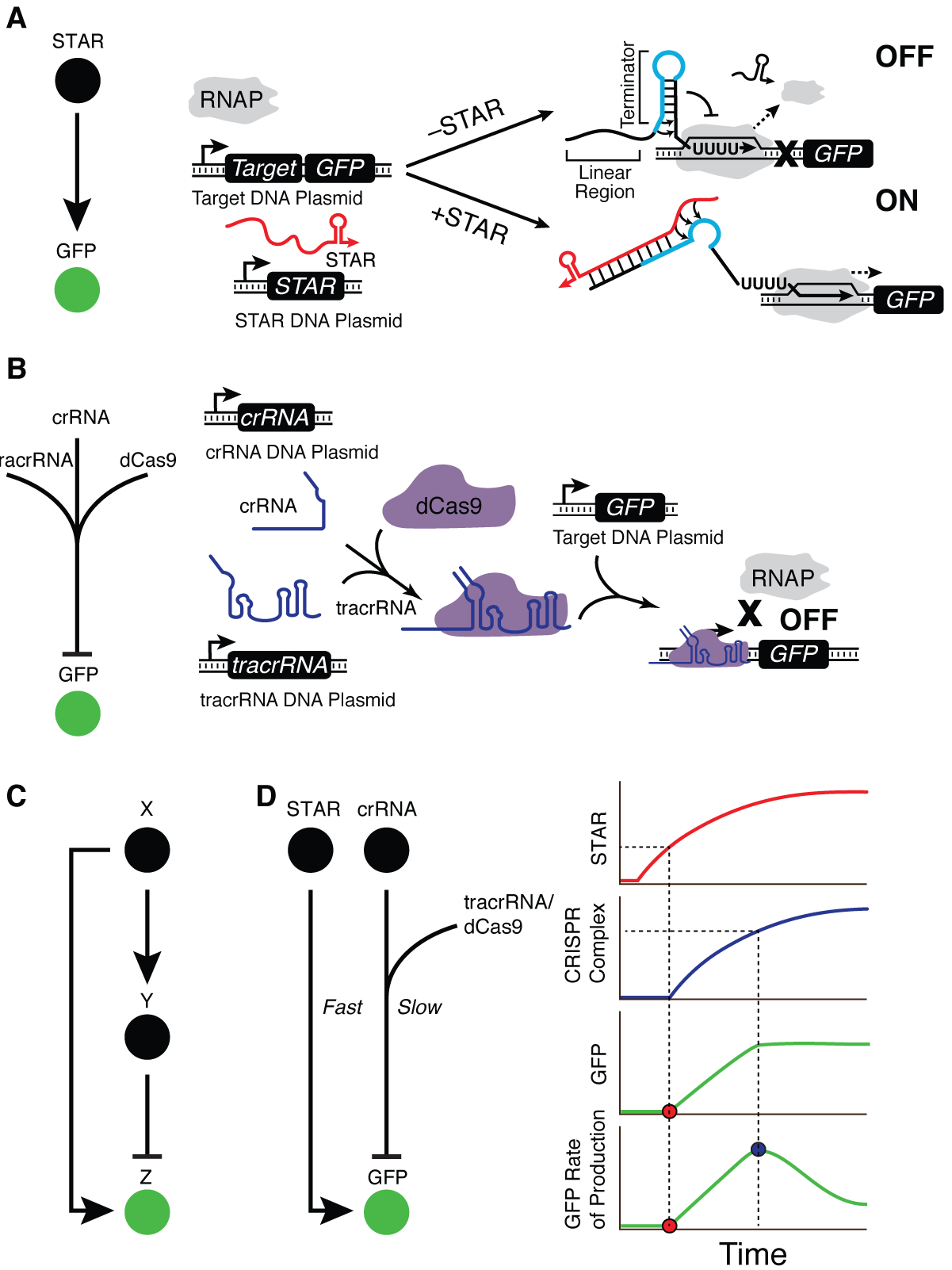
Architecture of a type 1 incoherent feedforward loop (I1-FFL) composed of STAR activation and CRISPRi repression. (A) Small transcription activating RNA (STAR) mechanism. The target RNA sequence folds into a transcriptional terminator (blue) that causes RNA polymerase to ratchet off the DNA complex and halt transcription upstream of the gene (gene OFF). When present, a STAR (red) binds to both the linear region and the 5’ half of the terminator hairpin (blue) of the target RNA, preventing terminator formation and allowing transcription elongation of the gene (gene ON). (B) CRISPR interference mechanism. The crRNA, tracrRNA, and dCas9 bind to form the CRISPR complex that specifically binds to a DNA sequence encoded by the crRNA sequence. When bound the CRISPR complex either blocks transcription initiation or transcription elongation. (C) The I1-FFL motif consists of three parts. An activator X activates expression of Z and its repressor, Y. (D) The pulse generator circuit works by taking advantage of fast STAR activation and slow CRISPRi repression. STAR activates GFP expression immediately while the crRNA/tracrRNA/dCas9 formation causes a delay before finally repressing GFP expression. In TXTL there is no protein degradation, so this causes a pulse in the rate of GFP production.

CRISPRi is a method of transcriptional repression that relies on targeting a catalytically dead Cas9 (dCas9) nuclease to a gene.^12^ Targeting is dictated by a guide RNA (gRNA) with a 20-bp segment complementarity to the sequence of interest. Here we use the *Streptococcus pyrogenes* Cas9 that targets sequences flanked by a 3’ NGG PAM. Binding of the dCas9:gRNA ribonucleoprotein complex to DNA can either block polymerase binding if the targeted region is near a promoter or halt transcription elongation if the targeted region is within a gene. Orthogonal gRNAs can be designed to independently regulate multiple genes or to integrate signals for genetic circuits such as logic gates.^14^ In nature, gRNAs are produced by RNase III cleavage of dsRNA formed by the binding of a trans-activating crRNA (tracrRNA) to complementary sequences in a transcribed CRISPR RNA.^15^ The resulting processed crRNA binds to Cas9 (or dCas9) to form an active ribonucleoprotein complex (Figure 1B). CRISPRi works efficiently using either gRNAs produced by the processing of crRNA/tracrRNA duplexes or using single-guide RNAs (sgRNAs) which fuse the tracrRNA and crRNA to mimic the processed form using a single molecule.^16^ In this work we use separate crRNA and tracrRNA because they represent the natural form of the gRNA as it is expressed in bacteria, and they also add to an additional time delay in the CRISPRi regulation due to the kinetics of pairing between the RNAs.

One difference between STAR and CRISPRi mechanisms is the timescale on which the regulation occurs. STARs rely on one co-transcriptional RNA-RNA interaction that results in transcription activation typically within minutes,^2^ while CRISPRi requires the formation of an RNA-protein repressor complex before DNA binding to DNA for repression, which has been shown to take on the order of one hour for regulation to occur.^12^ This timescale difference between these two opposing modes of gene regulation thus creates an intriguing possibility to use STARs and CRISPRi to engineer a network that produces a pulse of gene expression, similar to the incoherent type-1 feedforward loop (I1-FFL).^17^

The I1-FFL is a common network motif in natural bacterial networks^18-20^ and has received much interest due to its ability to produce a pulse of gene expression^17,21^ and accelerate the response time.^22^ I1-FFLs have also been used to implement band-pass filters,^23,24^ fold-change detection,^25^ biosensing,^26^ and noise buffering.^27^ An I1-FFL consists of an activator X that activates a gene Z and simultaneously its repressor, Y (Figure 1C). It can produce a pulse of gene Z expression because the activation reaction is triggered immediately by X, while the dominating repression occurs with a delay due to the presence of the intermediate component Y.^17^ Here, we exploit STARs to induce rapid activation of gene expression, and CRISPRi to achieve delayed repression due to the slow assembly of the gRNA-dCas9 complex. We expect that, when combined, these two RNA-based regulatory mechanisms will operate on timescales that are sufficiently different to yield a transient pulse of gene expression (Figure 1D). While our design is not an I1-FFL by a strict definition, it accomplishes the same general behaviour and should produce a pulse of gene expression by exploiting the regulatory timescale differences to cause the delayed repression of Z after fast activation.

A challenge in interconnecting molecular components characterized in isolation is that unexpected interactions between species and resource competition can affect the predicted operation of the composed system, as demonstrated previously.^28^ Reaction rates can be affected by possible crosstalk between the components and the relative abundance of RNA species and dCas9, which are subject to biological noise and circuit complexity,^29^ thus making the prediction of the integrated construct dynamics necessary and challenging. To address these challenges, we use an interdisciplinary approach that combines cell-free experiments and mathematical modeling.

Mathematical models have gained popularity in guiding the construction and characterization of dynamic molecular systems, given their cost-effectiveness and efficiency as compared to experiments.^30-33^ Ordinary differential equations (ODEs) are an effective tool to model molecular reaction networks, gene expression in protein-based genetic network systems,^18,34^ and small RNA transcriptional circuits.^33,35^ ODEs are particularly suitable to model and parameterize cell-free reactions, where initial concentration of chemical species can be accurately controlled. In order to rapidly characterize the STAR and CRISPRi reactions we developed ODE models based on experiments performed with TXTL, an *E. coli* cell-free transcription-translation platform.^36^ TXTL experiments have been successfully combined with mathematical models to parameterize and understand RNA circuits.^33,37,38^ TXTL is ideal for prototyping genetic circuit dynamics because it is quick and easy to use, requires minimal cloning, and shows good agreement with *in vivo* data,^39^ and recently it was used to to characterize CRISPR nucleases and guide RNAs.^40^ Additionally, TXTL also allows for experiments that would otherwise be difficult to perform *in vivo* by giving direct control over component concentrations and enabling circuit optimization and flexibility when designing experiments to fit model parameters.

Here, we start by using TXTL to verify that the STAR and CRISPRi present sufficiently distinct regulatory timescales. Then, we build ODE models for the STAR and CRISPRi pathways in isolation, and we perform systematic TXTL experiments to parameterize and validate the models. We find that when the models are composed to build the IFFL circuit, they predict the expected pulse generation. We conclude with experiments showing that, when connected together to regulate the same promoter, the candidate STAR-CRISPRi pulse generator circuit yields a pulse in target gene expression, and that the composed models can quantitatively capture the pulse generator behavior. Our results demonstrate that the combination of modeling and experiments in a simplified TXTL environment is an effective approach to prototyping biological dynamic circuits for control of gene expression. Most importantly, our results indicate that RNA regulators characterized in isolation can be combined in more complex circuits without loss of performance when interconnected, making them modular and composable components for dynamic synthetic circuits.

## RESULTS

### Pre-incubation experiments confirm the expected STAR/CRISPRi timescale difference

We first sought to verify the timescale difference between STAR and CRISPRi regulation expected from previous studies.^2,11^ To do this, we designed experiments that isolated the kinetic processes of each mechanism. We transcribed RNA components and allowed folding and complex formation with previously synthesized dCas9 before assessing regulatory function, to isolate only the timescale of the regulatory mechanism. When performing a typical TXTL experiment, all DNA is added to the reaction at t = 0 and gene expression is measured over the course of a few hours. Inherent to this experimental design is a delay due to the transcription of RNA regulator parts, which must first be transcribed before they can perform their function. In order to isolate the timescale of the regulatory event, we incubated a plasmid expressing each RNA regulatory part alone for 2 hours, essentially allowing the TXTL reaction to synthesize RNA regulators before being assessed for function. We then mixed pre-incubated reactions with reporter DNA and characterized the response time of the system. In this way, we removed the timescale needed for regulatory RNA synthesis and instead focused the characterization experiment on the relevant timescales of action for each regulator.

The STAR system only has one trans-acting RNA, so we incubated a plasmid expressing the STAR RNA or a plasmid expressing a non-functional control RNA in TXTL for 2 hours. We then added DNA encoding the p70a-Target-GFP plasmid to this reaction mixture at t = 0 and began measuring fluorescence over time. We observed detectable STAR activation of gene expression ~20 minutes after the addition of the GFP plasmid (Figure 2A) and STAR activation as determined by the GFP production rate reached 70% of the steady state after 35 minutes, where the steady state was computed from an exponential fit (Figure 2B) as described in Supplementary Note S1.

**Figure 2.**
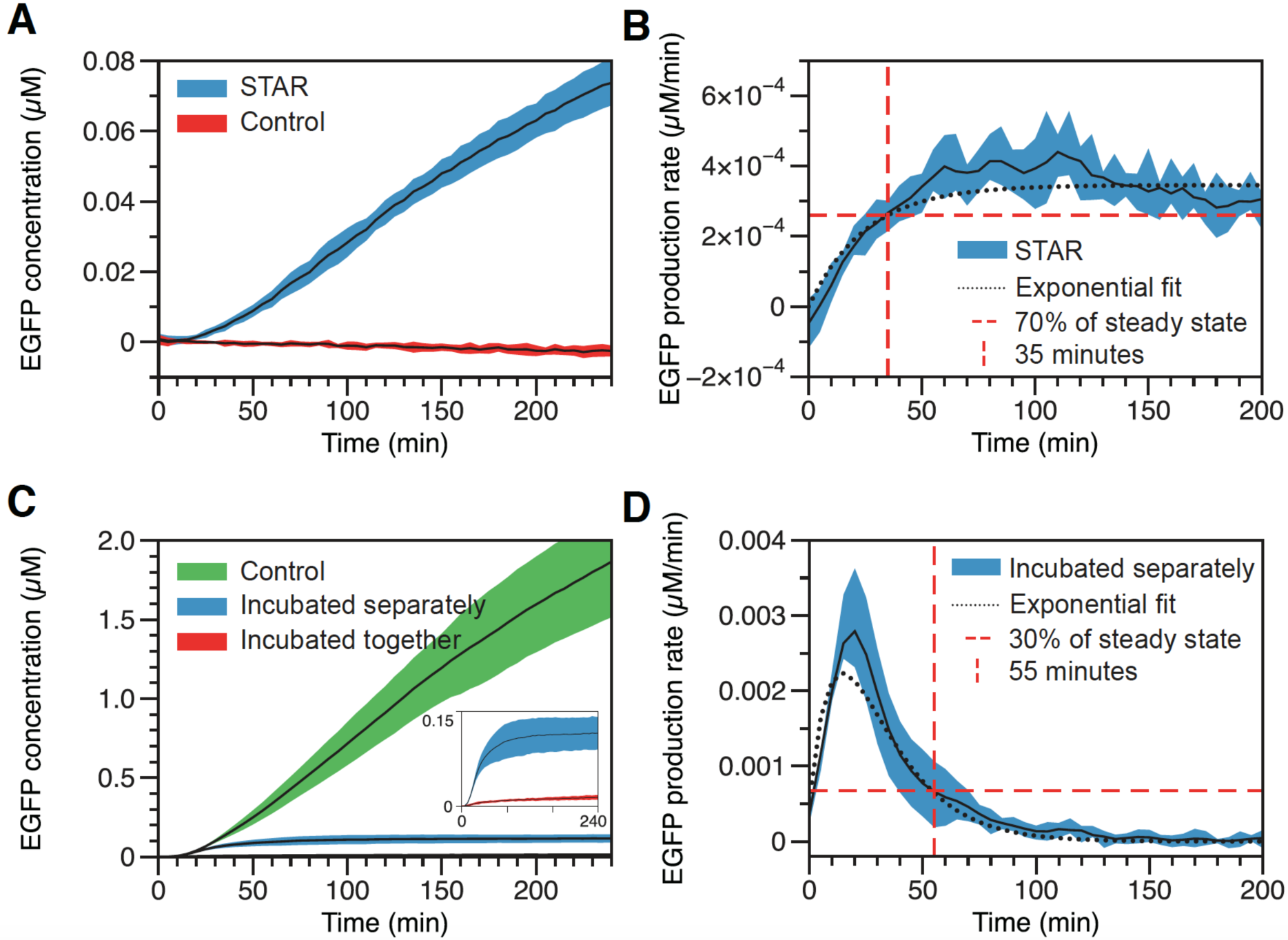
Pre-incubation experiments indicate that STAR activation is faster than dCas9-based repression. (A) Functional time course characterization of GFP expression when STAR is pre-incubated (blue) or a non-function control is pre-incubated (red). The timescale of STAR activation is on the order of 20 minutes after reporter DNA is added to the reaction. (B) The production rate of GFP expression for the STAR pre-incubation experiment (blue). The GFP production rate reached 70% of max as determined by the exponential fit (dotted black line) at 35 minutes. (C) Functional time course characterization of the CRISPRi response when parts are incubated together (red) or separately (blue) in comparison to unrepressed expression (green). The timescale of pre-incubated CRISPR repression is much faster than when the parts are incubated separately, suggesting that the dCas9 loading time adds a significant delay to the system. The inset shows the two repressed states. Data for all pre-incubation combinations of CRISPRi parts is shown in Supplementary Figure S6. (D) The production rate of GFP expression for the CRISPRi pre-incubation experiment (blue). The GFP production rate reached 30% of its peak as determined by the exponential fit to the derivative (dotted black line) at 55 minutes.

We anticipated that regulation of gene expression by the dCas9 complex would take significantly longer than the STAR activation, given previous observations suggesting that gRNA loading onto dCas9 takes on the order of ~1hr in the presence of non-specific RNAs.^41^ As the CRISPRi system requires a crRNA, tracrRNA, and dCas9, a more sophisticated experiment was required to characterize the regulatory timescale. Specifically, we sought to determine the timescale for crRNA-tracrRNA-dCas9 complex assembly required for the dCas9 complex to repress gene expression. To quantitatively estimate this timescale, we incubated the DNA encoding each RNA component in all combinations of alone, together, and in TXTL already containing dCas9 for 2 hours (Supplementary Figure S1) and then combined them into a final reaction with DNA encoding the p70a-GFP plasmid before began measurement. For clarity, we only show two conditions in Figure 2C: all alone or all together in TXTL containing dCas9. When incubated separately, we expect all components to be present at high concentrations at the beginning of the measurement but no CRISPRi repression complex will have formed yet. The complex will begin forming when the measurement starts. When incubated together, we expect the CRISPRi complex to have already formed and be present at high concentrations. Comparing these two conditions indicates the time it takes for the crRNA-tracrRNA-dCas9 complex to form and then repress (Figure 2C). However, when incubated separately, the complex was slower to repress gene expression, and did not achieve full gene repression until 55 minutes after addition of the DNA reporter construct (Figure 2D). This large difference in response times reveals that the crRNA-tracrRNA-dCas9 complex takes on the order of 55 minutes to fully form and perform its function in TXTL, which is similar to previous research.^41^

Taken together, these results indicate that there is a timescale difference between STAR activation (70% of the steady state production rate seen after 35 minutes) and CRISPRi repression (30% peak production rate seen after 55 minutes) due to the extra steps required for the crRNA-tracrRNA-dCas9 complex assembly as opposed to the direct RNA-RNA interactions of the STAR mechanism. These timescale differences could therefore be exploited to construct a simple network architecture that produces a pulse of gene expression.

### STAR and CRISPRi model derivation

After verifying the timescale difference with our pre-incubation experiments, we then sought to construct mathematical models for the STAR and the CRISPRi systems respectively, to computationally test our hypothesis and guide the design of the circuit, before conducting further experiments. We used ordinary differential equations to model the rate-of-change of each molecular concentration, as a result of coupled kinetic reactions (Figure 3).

**Figure 3.**
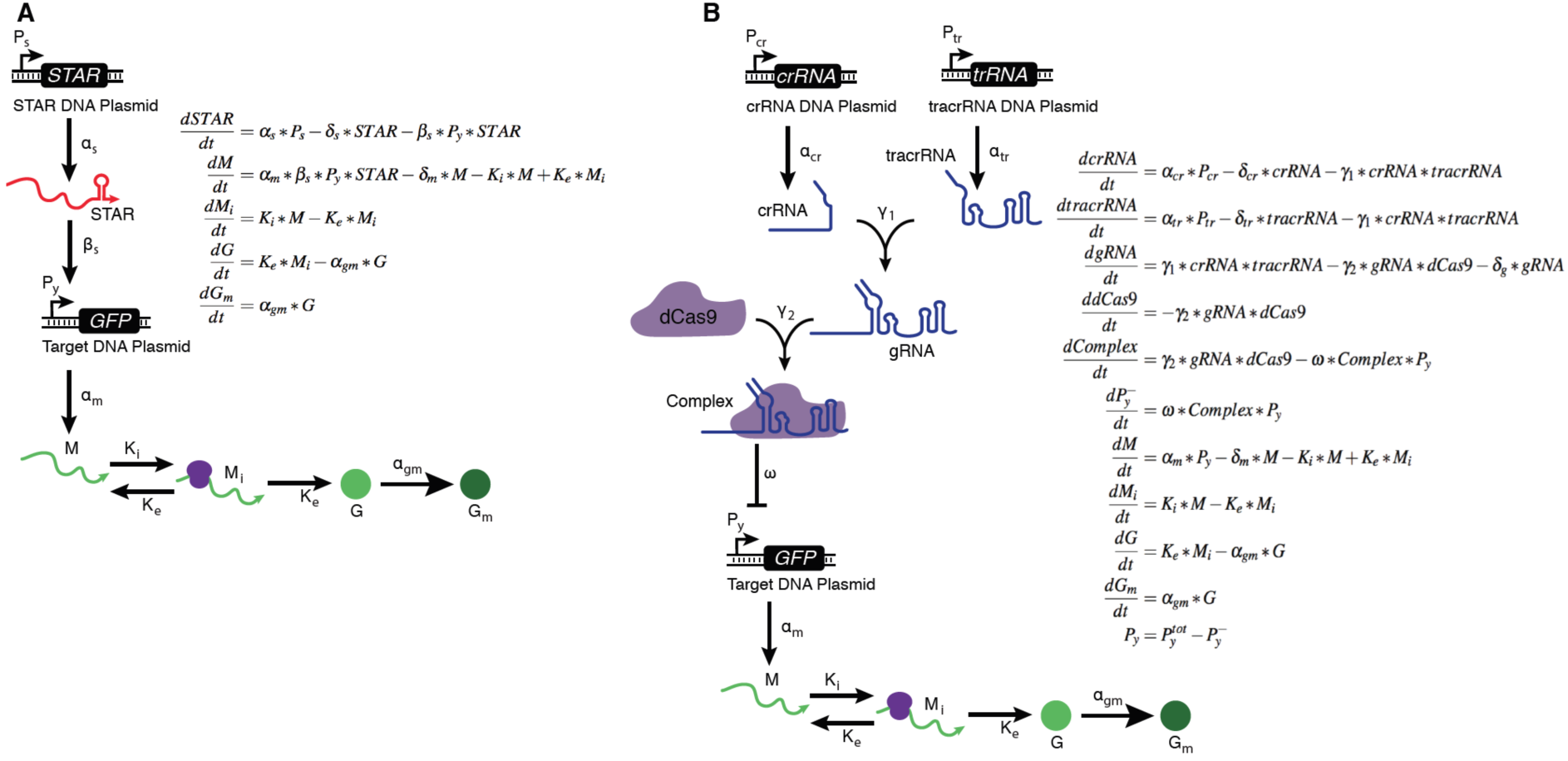
Separate STAR and CRISPRi models with the corresponding topology. The STAR activation is modelled as a one-step binding at rate β_s_, to the free output promoter P_y_, to enable expression of GFP mRNA M. The CRISPRi repression is modelled as a two-step reaction, where formation of active repressor complex happens before it binds to the free output promoter P_y_ to form the repressed P_y_^-^, and GFP is only expressed from the free P_y_ promoter. For simplicity, the degradation rates of the RNA species are modelled but not shown in the topology. In both models, mature GFP protein G_m_ is compared to experimental measurements. All the STAR, M, Mi, G, G_m_, crRNA, tracrRNA, gRNA, Complex, and P_y_^-^ are initiated with concentration 0 nM. The initial free P_y_ plasmid was 0.5 nM, and dCas9 concentration was estimated to be 35 nM based on previous experimental measurement.

We modelled the STAR activation as a one-step reaction, where STAR binds to the free promoter P_y_ directly to achieve transcription activation, at rate β_s_, mimicking the fact that STAR activation only requires RNA-RNA interactions. This is an approximation that coarse-grains the details of how the small RNA modifies target RNA structure to activate transcription, but it is justified based on similar simplifying assumptions made in previous work modeling RNA transcriptional repressors.^33,37^ In parallel, we modelled CRISPR-Cas9 complex formation as a two-step reaction process. As part of this process, the tracrRNA and the crRNA bind to form the gRNA at rate γ_1_. The gRNA can then bind to dCas9 to form the active repressor complex at rate γ_2_. Since dCas9 dissociation rates are extremely low with no mismatches,^42^ we assumed the formation of the CRISPR-Cas9 complex and its binding (at rate ω) to the free promoter P_y_ to form the repressed promoter P_y_^-^ to be irreversible. While capturing the key reactions in the CRISPR-Cas9 formation, the model coarse-grains the detailed dynamics of how crRNA, tracrRNA and dCas9 interact with each other and interferes transcription. To enable a direct comparison between the STAR and CRISPRi regulation pathway, we used a first-order kinetic reaction to model the STAR activation, instead of the Hill-type function used in Hu et al.^33^

In the STAR system, reporter p70a-Target-GFP mRNA (M) is only produced when p70a-Target-GFP (P_y_) is activated (i.e. bound to STAR at rate β_s_), at rate α_m_, while in the CRISPRi system, M is only produced from the free promoter p70a-GFP (for simplicity and for later use in the combined model, this is also denoted by P_y_), at rate α_m_. The GFP translational initiation, elongation, and maturation were modelled following prior work,^33^ and the mature GFP (G_m_) is compared to the experimental measurement.

In addition to the transcriptional rates above, each RNA species has a degradation rate and each protein species has a translation rate. Specifically, α_s_, α_cr_, α_tr_, δ_s_, δ_cr_, δ_tr_, and δ_g_ are the transcriptional and degradation rates of STAR, crRNA, tracrRNA, and gRNA respectively; δ_m_ is the degradation rate of GFP mRNA, M; K_i_ is the translation initiation rate, K_e_ is the translation elongation rate, and α_gm_ is the GFP maturation rate. P_y_^tot^ is the total amount of reporter promoters, M_i_ is the translationally initialized mRNA, and G is the immature GFP protein. We note that no protein degradation rate is included because proteins do not degrade in TXTL unless degradation tags are included,^43^ and there is no translation rate for dCas9 because extracts were made from *E. coli* cells expressing dCas9.^40^

### Model parameterization

With the separate STAR and CRISPRi models, our next step was to extract suitable kinetic parameters to construct a combined model for reliable predictions. To achieve this, we adopted a Bayesian inference parameterization approach^44,45^ to fit parameters for STARs and CRISPRi separately. Specifically, we used three sets of the STAR activation experiments (Full experimental data shown in Supplementary Figure S2) to train our model for the three STAR-related kinetic parameters: α_s_, δ_s_, and β_s_. We also used three sets of the CRISPRi repression experiments (Full experimental data is shown in Supplementary Figure S3) to train our model for the eight CRISPRi-related kinetic parameters: α_cr_, α_tr_, δ_cr_, δ_tr_, δ_g_, γ_1_, γ_2_, and ω, As crRNA and tracrRNA were transcribed from the same promoter in our experiments, we assumed that they share the same transcription rate, and we set α_cr_ = α_tr_ in the fitting. The five reporter GFP-related parameters (α_m_, δ_m_, K_i_, K_e_, and α_mg_) were also fitted for both STAR and CRISPRi.

For both STAR and CRISPRi experiments, we initiated our fitting with 10 different initial guesses that were evenly spaced in the admissible parameter intervals that were inferred from previous publications (Supplementary Table S1).^33^ To fit the 8 parameters in the STAR model, we conducted 105,000 iterations of parameter updates to seek convergence, and to fit the 12 parameters in the CRISPRi model, we conducted 210,000 iterations. The probability of accepting parameter set *i* from parameter set *j* was set according to the following:^44^

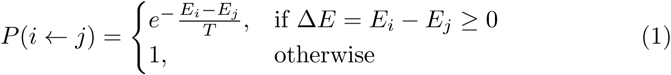

with T = 0.125 (= 2σ^2^), and σ is the estimated measurement error. The cost function *E* is defined as the cumulated point-wise squared prediction-measurement error for each experiment cycle:

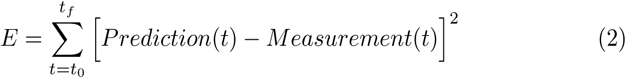

The parameter set that gave the lowest cost function *E* across all the fittings was deemed as the best fitted parameter set. The corresponding simulations are plotted in Figure 4A and B against the experimental measurement. The comparisons between predictions and data demonstrate that models trained with the Bayesian inference approach were able to reproduce the dynamics of the STAR and the CRISPRi system under various conditions. To understand the distribution of each parameter, we ranked all the sampled parameter sets (i.e. 10 x 105,000 and 210,000 sets of parameters for the STAR and CRISPRi fitting respectively) with respect to the corresponding value of the cost function *E*. Figure 4C shows the parameter distribution of the first 1000 sets of parameters that gave the lowest fitting error *E*. Note that the five GFP-related parameters shown in Figure 4C were fitted from the STAR activation experiments, for demonstration. The values of the best fitting parameters are given in Supplementary Table S1.

**Figure 4.**
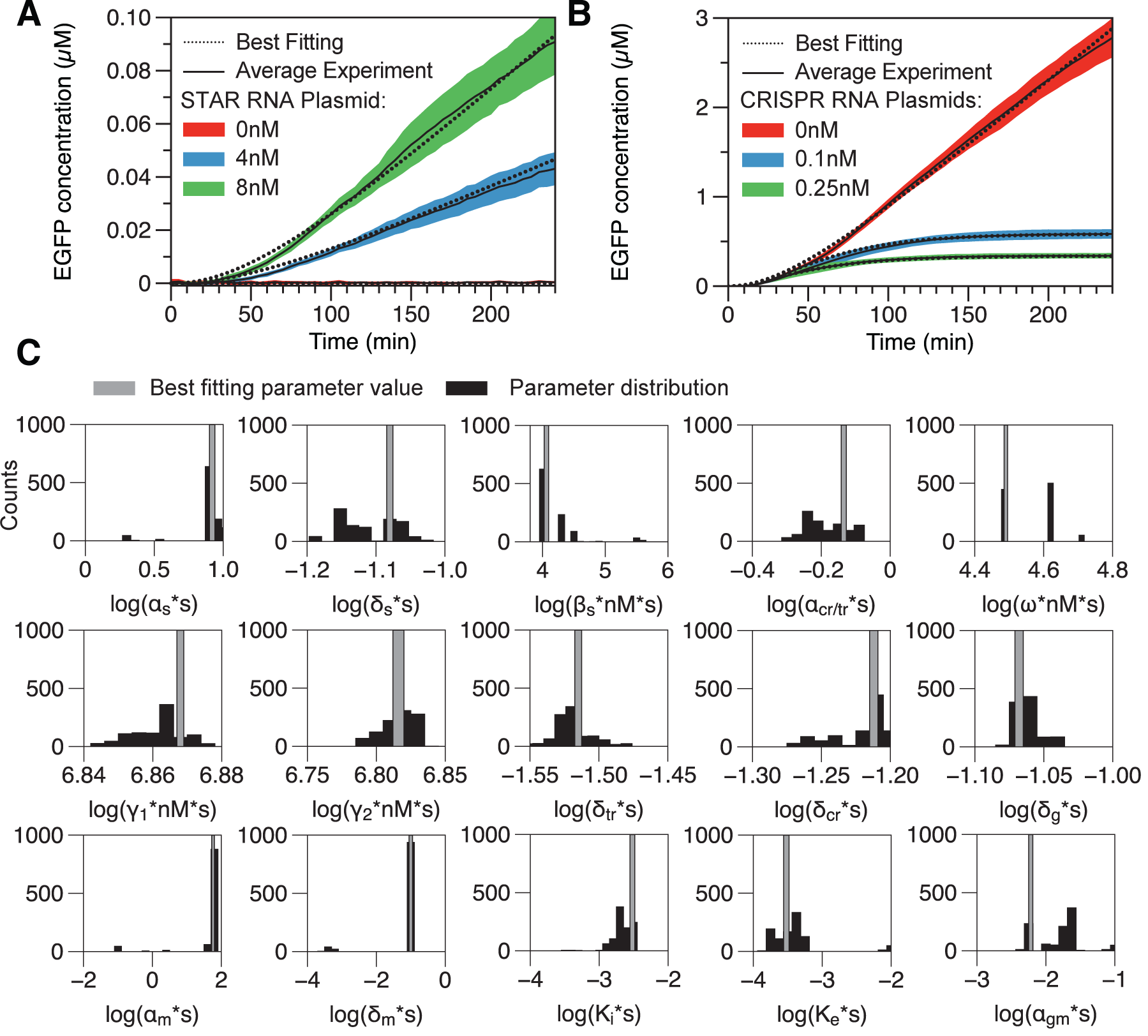
Model parameterization with separate STAR and CRISPRi experiments. (A) Comparison of best fitted simulation to the STAR experiments for three conditions: high activation with 8 nM of STAR plasmid (green plots, 8 nM STAR), moderate activation with 4 nM of STAR plasmid (blue plots, 4 nM STAR), and no activation with no STAR plasmid (red plots, STAR OFF). (B) Comparison of the best-fitted simulation to the CRISPRi parameterization experiments for three conditions: no repression with no crRNA or tracrRNA (red plots, 0nM CRISPR RNA), moderate repression with 0.1 nM crRNA and tracrRNA plasmid (blue plots, 0.1 nM CRISPRi RNA), and complete repression with 0.25 nM crRNA and tracrRNA plasmid (green plots, 0.25 CRISPRi RNA). (C) Histogram of parameters obtained from 1000 samples that gave the lowest fitting error within the pool of 10 x 105000 and 10 x 210000 fitting rounds for the STAR and CRISPRi system respectively. Grey bar indicates the location of the parameter value that gave the best fitting. Note, all the kinetic parameters are scaled to be dimensionless before taking their log values in the histogram plots.

Interestingly, while some CRISPRi-related parameters have a relatively wide distribution, we see limited variation in the repressor formation-related parameters such as ω for the plotted 1,000 fitted parameter values. This observation suggests that the repressor formation kinetics dominate the accuracy of the CRISPRi regulation process. On the other hand, all three STAR-related parameters displayed a relatively wide distribution, which suggests the existence of multiple optimal solutions for the fitting. This might be due to our simplification of the STAR activation mechanism and/or limited experimental conditions (e.g. initial concentrations), such that a wide range of parameter values can fit well the model. Note that fewer reaction steps and experimental conditions lead to fewer constraints for the parameterization. The corresponding fitted parameter distribution and correlation are given in Supplementary Figures S4 and S5.

### Pulse generator modeling and experimental verification

After parameterizing the separate STAR and CRISPRi models, we then combined them to build the pulse generator model by introducing a competition for P_y_ promoter binding between STAR and CRISPRi (Figure 5). In the combined pulse generator model, a free promoter P_y_ can either bind to CRISPRi to form a repressed state or to STAR to form an activated state for gene expression. Once P_y_ is bound to the CRISPRi complex, it becomes unavailable for STAR activation. To simulate the model, all output promoter copies were initiated in the free state (unbound), with a fixed concentration to mimic conditions used in the CRISPRi characterization experiments.

**Figure 5.**
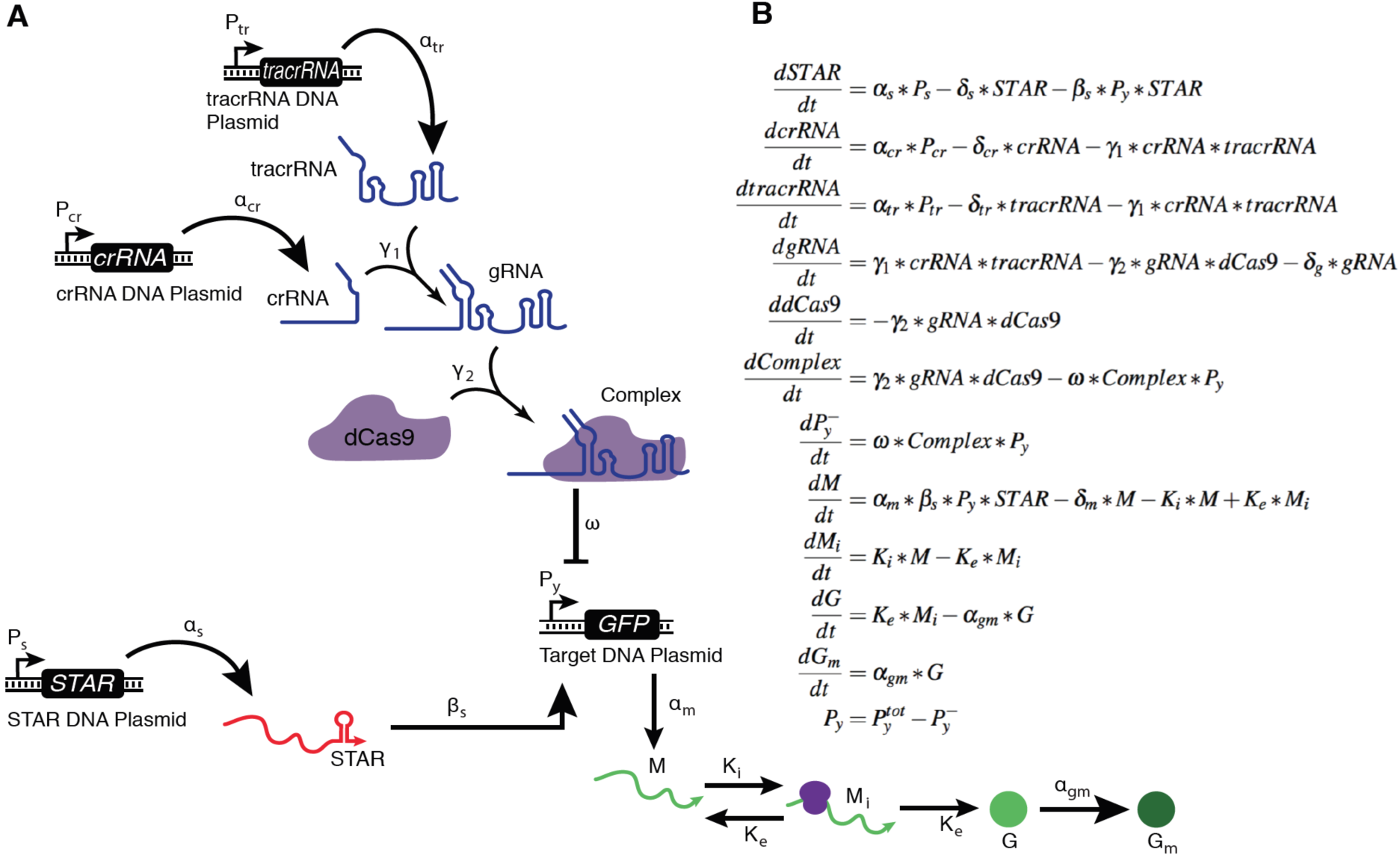
Topology of the pulse generator model. The separate STAR and CRISPRi model are combined by introducing a competition for P_y_ binding through the repressor formation in dP_y^-^_/dt and the activation in dM/dt equations. Once CRISPRi repressor complex binds to P_y_ to form repressed state P_y_^-^, it can no longer be activated for expression. P_y_ and P_y_^-^ follows mass balance with a total initial concentration of P_y_^tot^. For simplicity, the degradation rates of the RNA species are modelled but not shown in the topology. All the STAR, M, Mi, G, G_m_, crRNA, tracrRNA, gRNA, Complex, and P_y_^-^ are initiated with concentration 0 nM. The initial free P_y_ plasmid was 0.5 nM, and the dCas9 concentration was estimated to be 35 nM based on previous experimental measurement.

We then used the combined model to test if a pulse could be generated in the production rate of the target gene. Instead of using one best fitted parameter set, we decided to combine the set of best fit from each of the 10 Bayesian fittings for both the STAR and the CRISPRi regulator experiments, obtaining 100 sets of parameters (10 STAR x 10 CRISPRi) to generate predictions of the pulse generator behaviour. This is because a mismatch between our model prediction and the pulse generator behaviour could be caused by the fact that multiple optimal parameters exist for each individual regulator model (Figure 4), so the combination of the very best fits might not give the most accurate prediction for the interconnected circuit. The procedure to generate combinations of best fits is summarized in Figure 6A. Prediction with the best STAR and CRISPRi separately fitted parameters demonstrated a plateau in the GFP concentration (dashed black plot in Figure 6B), and a pulse in the production rate (dashed black plot in Figure 6C), and indeed all 100 parameter combinations suggested a pulse in the production rate (Supplementary Figure S6). Given these observations, we expect the integrated pulse generator to function robustly and also to produce a pulse in experiments.

**Figure 6.**
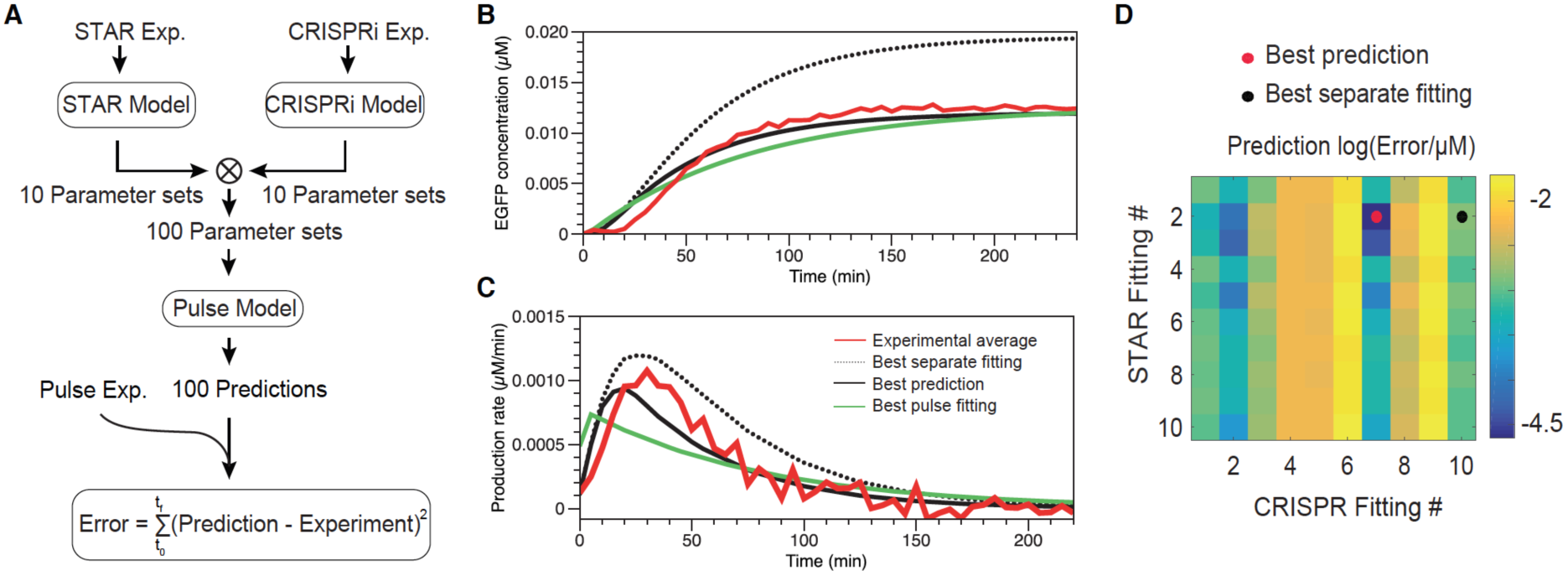
Pulse model prediction and experimental verification. (A) Procedure for parameterization and prediction: each of the individual STAR and CRISPRi models were trained with experimental measurements to fit 10 sets of best fitting parameters. These parameters were then combined into 100 sets that were used to predict the dynamics of the pulse model. The predictions were then compared to experimental measurement and quantified by the squared error between the prediction and the observed trajectories. (B) GFP concentration reached steady state in both the simulation (black and green) and experiments (red) within 240 min, while the best fitted parameter set predicted a higher steady state concentration level (dashed black), and the best prediction from the separately fitted parameters (solid black) gave better accuracy to the best fitting of the pulse model (green); (C) GFP production rate demonstrated a pulse which peaked at around 40 min and dropped when the repression kicked in and RNA degradation took over, in both the simulations (black and green) and the experiments (red). (D) Presentation of the prediction accuracy with the 100 sets of separately fitted parameters indicates the best separately fitted parameter set did not give the best prediction in the combined pulse model. Red dot indicates the location of the best separately fitted parameter sets and the yellow dot indicates the location of the parameter set for the best prediction. Note that they are in the same row (i.e. same STAR fitting trial) but different columns (i.e. different CRISPR fitting trial). Best separate fitting: prediction with parameters that best fit the STAR and CRISPR system individually; best prediction: the best of the 100 predictions with individually fitted parameters; best pulse fitting: best out of the 10 fittings to the pulse experiments.

We then performed a TXTL experiment that combined both the STAR and the CRISPRi systems. As in the separate CRISPRi experiment, we added 0.25 nM crRNA, 0.25 nM tracrRNA, and 0.5 nM p70a-Target-GFP. Since the STAR ON expression level is significantly lower than that of the CRISPRi system (Figure 4), we doubled the amount of STAR plasmid used in the separate STAR experiment from 8 nM to 16 nM in the combined system, to mitigate this difference. After the addition of all DNAs, we immediately began measuring fluorescent GFP expression (see Supplementary Figure S7 for complete experimental data). As predicted, the experiments also demonstrated a plateau in GFP expression level (Figure 6B, red), and a pulse in the production rate (Figure 6C, red). We only see a pulse in the production rate because TXTL has negligible protein degradation.^36^ If performed *in vivo*, we would expect a pulse in concentration rather than production rate. We then quantified the prediction accuracy by defining the prediction error in the same way as the cost function in Eqn. 2 to study the possible changes in the model parameters caused by the combination. The log-based prediction errors are summarized in the heat map in Figure 6D.

One interesting observation is that the best prediction (solid black plot in Figure 6B and C) was not achieved by the set of the best-fitted parameters (dashed black plot). The best-fitted parameter set predicted a higher steady-state concentration in GFP and a taller pulse in the production rate, as compared to the averaged experimental measurement (solid red plot in Figure 6B and C) and the best prediction. Indeed, the best fitted and the best prediction parameters were from the same STAR (same row in Figure 6D) but a different CRISPRi fitting trial (different column in Figure 6D). The values of the best prediction parameters are given in Supplementary Table S1. This observation suggested that the coupling may affect the CRISPRi dynamics, such that the set of parameters fit best the separate experiments but underpredicted the repressor formation rate, which lead to a higher predicted GFP expression level. A detailed parameter-to-parameter comparison between the best prediction and the best-fitted parameters is given in Supplementary Figure S8, to visualize the relative location of each parameter value.

We next asked how well we can fit the STAR/CRISPRi combined model to the experimental measurements, and how that compares to the best prediction with the separately fitted parameters. Again, to seek convergence we conducted 10 Bayesian fittings from different initial guesses, with 210,000 iterations for each fitting (same as in the CRISPRi fitting). The fitting that yielded the lowest fitting error is plotted in Figure 6B and C in green. Surprisingly, the best prediction with the separately fitted parameters slightly outperformed the best fits on the combined model. This could be due to the fact that in the combined model more parameters have to be simultaneously fitted relative to the individual component models, leading the combined model to require a larger number of samplings (i.e. initial guesses and/or iterations) to reach an equally good fit. Indeed, the fitting error comparison in Supplementary Figure S9 suggests that to fit 12 parameters in the CRISPRi model, even more iterations might be needed. Additionally, the best prediction from the separately fitted parameters is similar to the best fits out of 100 fittings, since it is the best prediction from a 10 x 10 best fitted parameter sets. To improve the fitting on the combined model, one can use more initial guesses and increase the number of iterations. Supplementary Figure S9 summarizes the detailed comparison of the accuracy and the error convergence for the STAR, CRISPRi, and pulse generator model fitting, respectively.

### DISCUSSION

In this work we have demonstrated an RNA-based pulse generator in TXTL that harnesses the difference in speed between STAR and CRISPRi regulation. This STAR-CRISPRi hybrid construct is able to produce a pulse of gene expression. STAR activation involves a single, fast, co-transcriptional RNA-RNA interaction while CRISPRi requires the slow formation of an RNA-protein complex leading to a delay before CRISPRi repression sets in. Combined, these mechanisms produce pulse of gene expression caused by the transcription of a few RNA molecules.

There have been a number of synthetic I1-FFLs built using protein regulators.^23,26,46^ Recently, we built an RNA-based I1-FFL that uses AHL to activate expression of a STAR RNA that activates expression of mRFP as well as a gRNA and dCas9 that repress mRFP.^2^ This design relies on an additional RNA cleavage strategy, cascading RNA regulatory events, and slow dCas9 production. Here, we constructed a simpler network that implements the pulse of gene expression of an I1-FFL, but faster and more effectively with a simpler network design.

As synthetic networks grow in complexity, models will be vital for predicting their behaviour and understanding dynamics, as they provide faster assessments of the network as compared to experiments. Here, we constructed and parameterized a coarse-grained mechanistic model and used it to predict the dynamics of the pulse generator network. With the simulation results, we observed possible modularity of the STAR regulator when combined with other structures to form more complicated networks, while the performance of the CRISPRi regulation might be affected, as indicated by the change in the parameter values. However, this observed change in the CRISPRi regulation might be due to several reasons: first, given the limited amount of training data, it could be possible that the CRISPRi parameters were over-fitted on the training data thus giving a non-ideal prediction in the new condition (combined system). Indeed, the complexity, parameterization methods, and experimental noise could all contribute to the accuracy of the model parameterization. Second, un-modelled (and undesired) coupling of the two regulatory pathways could affect the dynamics; for example, indirect competition for the transcription machinery could reduce transcription rates in a non-homogeneous manner in the two circuits, altering their regulation timescale. Third, the mechanism of the CRISPRi repressor formation might be oversimplified such that intermediate reaction steps were overlooked. For further investigations, we suggest a richer data set under various conditions for model parameterization, and a refined model to encompass more detailed reactions in the system.

Model parameterization can be challenging, especially when obtaining large amount of experimental measurements under various conditions is costly and a stochastic parameterization method is used, which would normally require convergence. The results in this study suggest that, instead of fitting all the parameters simultaneously, fitting part of a combined network separately could also lead to reliable predictions of an integrated structure, especially when the modularity of each component can be maintained. Because fitting parameters of individual modules for use in integrated structures provides a more computationally tractable alternative to comprehensive parameter fitting, we expect this approach to become predominant as synthetic molecular systems become more and more complex.

In summary, we demonstrate a STAR-CRISPRi hybrid pulse generator both with simulation and *in vitro* TXTL experiments; the circuit mimics the architecture and performance of an I1-FFL. We also demonstrated how mathematical modeling can be used to guide and assess the design of biological constructs. We found that parameters fitted from separate models can also accurately predict the performance of the combined model/construct. We further discussed the importance of sample data, and optimization settings in improving the parametrization. We anticipate the results in this study to provide guideline for future work in the modeling, parameterization, and construction of biological parts made of both STAR and CRISPRi regulators.

## MATERIAL AND METHODS

### Plasmid construction and purification

Key sequences can be found in Supplementary Table S2. All the plasmids used in this study can be found in Supplementary Table S3. The STAR plasmid and control plasmid were construct pJBL4971 and pJBL002, respectively, from Chappell et al.^2^ The GFP expression plasmid was p70a-GFP from Garamella et al.^43^ and the STAR-target plasmid was modified from this plasmid using iPCR. The plasmids expressing crRNA, tracrRNA, and the scrambled crRNA were constructed using Gibson Assembly and iPCR and sequence verified using sanger sequencing. Plasmids were purified using a Qiagen QIAfilter Plasmid Midi Kit (Catalog number: 12243) followed by isopropanol precipitation and eluted with double distilled water.

### TXTL Extract and Buffer Preparation

Cell extract and reaction buffer were prepared according to prior work.^43^

### TXTL experiments

TXTL buffer and extract tubes were thawed on ice for approximately 20 min. Separate reaction tubes were prepared with combinations of DNA representing a given circuit condition. Appropriate volumes of DNA, buffer, and extract were calculated using a custom spreadsheet developed by Sun et al.^36^ and modified to fit the experiments. Buffer and extract were mixed together and then added to each tube of DNA according to the previously published protocol. Each TXTL reaction mixture (10 μL each) was transferred to a 384-well plate (Nunc 142761), covered with a plate seal (Nunc 232701), and placed on a Biotek Synergy H1m plate reader. We note that special care is needed when pipetting to avoid air bubbles, which can interfere with fluorescence measurements. Temperature was controlled at 29°C. GFP fluorescence was measured (485 nm excitation, 520 emission) every 5 min. A calibration to EGFP concentration (μM) was performed using a standard curve of pure EGFP (Cell Biolabs STA-201) in order to present measurement data in terms of GFP concentration. Pre-incubation experiments were performed by combining two types of extracts. One extract has dCas9 pre-expressed and the other does not. Each plasmid was incubated in the appropriate extract and buffer for 2 hours before the pre-incubated reactions were combined in equal parts and measurements began.

### Modeling

Equations in Figure 3 were solved with MATLAB_R2014b ode23s solver to get the simulated GFP concentration for the error calculation (Eqn. 2) in data fitting. Candidate parameters were generated with a uniform distribution within a bounded interval (Supplementary Table S1), using MATLAB random number generation function *rand*. One trial of the Bayesian inference data fitting (i.e. one initial guess, with 105000 iterations) took about three computational hours on a Macbook Pro with a 2 GHz Intel Core i7 processor. Model in Figure 5 was also numerically solved with MATLAB_R2014b ode23s function to get predictions for the combined pulse generator.

## ACKNOWLEDGMENTS

This work was supported by the Defense Advanced Research Projects Agency (contract HR0011-16-C-01-34). The authors declare no conflict of interest.

## SUPPORTING INFORMATION

Supplementary information on detailed experiments, modelling, and data analysis.

